# Non-destructive *in situ* monitoring of structural changes of 3D tumor spheroids during the formation, migration, and fusion process

**DOI:** 10.1101/2024.07.25.605064

**Authors:** Ke Ning, Yuanyuan Xie, Lingke Feng, Wen Sun, Can Fang, Rong Pan, Yan Li, Ling Yu

## Abstract

For traditional laboratory microscopy observation, the multi-dimensional, real-time, *in situ* observation of three-dimensional (3D) tumor spheroids has always been the pain point in cell spheroid observation. In this study, we designed a side-view observation petri dish/device that reflects light, enabling *in situ* observation of the 3D morphology of cell spheroids using conventional inverted laboratory microscopes. We used a 3D-printed handle and frame to support a first surface mirror, positioning the device within a cell culture petri dish to image cell spheroid samples. The imaging conditions, such as the distance between the mirror and the 3D spheroids, the light source, and the impact of the culture medium, were systematically studied to validate the *in-situ* side-view observation. The results proved that placing the surface mirror adjacent to the spheroids enables non-destructive *in situ* real-time tracking of tumor spheroid formation, migration, and fusion dynamics. The correlation between spheroid thickness and dark core appearance under light microscopy and the therapeutic effects of chemotherapy doxorubicin and Natural Killer cells on spheroids’s 3D structure was investigated.

## Introduction

Three-dimensional (3D) tumor spheroids have emerged as a powerful tool for studying tumor biology and drug response. These multicellular aggregates better recapitulate solid tumors’ complex microenvironment and heterogeneity than traditional 2D cell culture models (R et al., 2014). 3D tumor spheroids exhibit gradients in oxygen, nutrients, and metabolites that mimic the pathophysiology of native tumors (Derda et al., 2009; Huh et al., 2011), making them valuable for investigating tumor growth, invasion, and response to therapies.

Visually, spheroids appear as translucent balls with a well-defined boundary and darker core when imaged by brightfield or phase contrast microscopy (Amaral et al., 2017; Pan et al., 2023b, 2023a). Spheroid growth is characterized by an initial exponential phase followed by a plateau once the spheroid exceeds a critical diameter (typically around 500 μm)(Amaral et al., 2017; Kim et al., 2021; Klowss et al., 2022). Thickness measurements can detect this transition and provide a more sensitive readout of growth arrest than the length in the *x* and *y* directions (Browning et al., 2021). Characterizing the morphology and volume of 3D tumor spheroids is essential for understanding their growth kinetics and evaluating treatment effects. Directly measuring width/diameter and thickness enables more precise volume calculations that could compensate for spheroids’ non-uniform shape and size (Napolitano et al., 2007).

However, light microscopy itself has limitations in quantifying the complex 3D structure of spheroids, especially the width and thickness of the spheroids simultaneously. The translucent nature of spheroids leads to blurred images that can cause overestimation of size and volume based on 2D measurements alone (Costa et al., 2019). Confocal imaging enables optical sectioning of spheroids to generate 3D reconstructions, and fluorescent labeling of cells allows direct measurement of thickness and diameter from cross-sectional views (Raza et al., 2020). However, light scattering and absorption limit imaging depth, making it challenging to resolve the core of larger spheroids (>300 μm)(Fang et al., 2023; Pan et al., 2023a; Steinberg et al., 2020). Optical coherence tomography (OCT) is a non-invasive imaging technique that uses low-coherence light to create cross-sectional images of tissue structure, allowing measurement of thickness and diameter even in larger spheroids (El-Sadek et al., 2021, 2020; I et al., 2020). Light sheet fluorescence microscopy (LSFM) is another emerging technique that illuminates samples with a thin sheet of light, enabling rapid 3D imaging of entire spheroids and measuring thickness and diameter from 3D reconstructions (Eismann et al., 2020; Paiè et al., 2023; Shi et al., 2024). However, LSFM requires specialized equipment and may be limited by light scattering in dense spheroid cores (Andilla et al., 2017; Costa et al., 2019).

Another challenge in the field is the lack of standardized protocols for measuring spheroid thickness and width across different imaging modalities and analysis software (Froehlich et al., 2016; Koudan et al., 2020; Moraes, 2020). Balancing spheroids thickness analysis while further observing changes in the sample’s three-dimensional structure is extremely challenging. Due to limited perspectives, current research tends to simplify the relatively regular morphology of spheroids into models that are centrally symmetrical or axis-symmetrical (Senavirathna et al., 2013). However, in addition to regular cell spheroid morphology, cell spheroids can undergo irregular changes. This inconsistency makes it difficult to compare results across studies and reproduce findings. Establishing guidelines for image acquisition, processing, and reporting is critical for advancing the field.

In this work, using conventional brightfield microscopy, we propose a side-view observation device to systematically study the correlation between bottom-view and side-view of 3D tumor spheroids. We designate the bottom view, visible through inverted microscope imaging, as the underside of the spheroid sample (*x-y* plane), where measurements such as width and diameter can be obtained. Images captured through the side-view observation device represent the side-view of the sample (*x-z* plane), from which measurements such as width, height/thickness, and contact angle with the *x-y* plane can be derived. First, the 3D spheroids side-view observation device was fabricated by assembling a cell petri dish, an agarose-microwell array, and an optical module for observing. The device was applied to track the dynamics of 3D cell spheroid formation and cell migration from the spheroid. In addition, the two spheroids’ fusion process and corresponding morphological changes at the bottom- and side-view were characterized to investigate the fusion dynamics. With the multi-angle observation view, the correlation between the thickness of the spheroid and the dark core appearance under the light microscope examination was studied. Last, the killing effect of chemotherapy compound Doxorubicin (DOX) and immune cell Natural killer cells (NKs) on the 3D structure of the spheroids were studied based on the multi-angle observation device.

## Results

### Validation of side-view observation-achieved by placing a first surface mirror adjacent to the 3D spheroids

The 3D spheroids were cultivated in agarose micro-wells within a standard petri dish. The assembled side-view observation device, featuring a 3D-printed handle and frame with an attached first surface mirror, was placed directly on the microscope stage (**Fig. 1A**). The magnets attract each other up and down the petri dish lid. Magnet pairs embedded in the handle and frame allow flexible movement, enabling the mirror to be positioned near the sample by moving the handle along the petri dish lid (**Fig. 1B**).

**Fig. 1.**
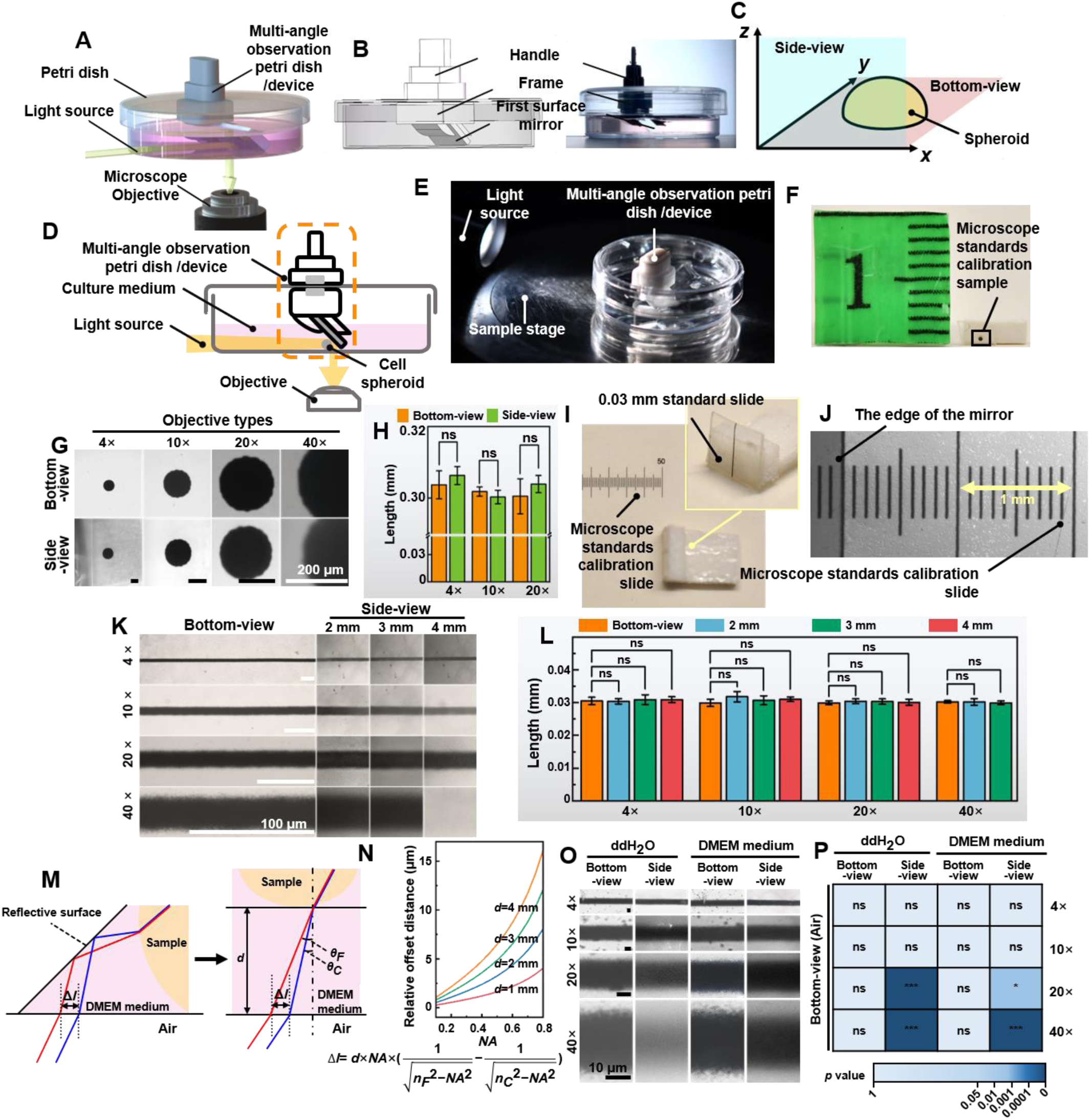
Validation of the non-destructive *in situ* observation of 3D spheroids by the multi-angel observation device. A) The illustration of the multi-angle observation device; B) Exploded view diagram of the multi-angle observation device; C) The definition of the bottom-view and the side-view of a spheroid sample; D) The working principle of the device; E) The photo of the petri dish mounted on the sample stage of the microscope. F) The 0.3 mm round-type microscope calibration slide for side-view observation. G) The bottom-view and side-view images of the 0.03 mm microscope calibration slide were captured with different objective lenses. H) The length of the 0.3 mm dot measured from the bottom-view and side-view captured images. I) A 0.03 mm line-type microscope calibration slide was vertically fixed to a base positioned at a 90° angle to the start of another microscope calibration slide. J) The photograph of the microscope calibration slide and the image of the edge of the first surface mirror (gray part) by the microscope. K) The images were captured with the distance between the mirror and the sample at 2 mm, 3 mm, and 4 mm. L) The length of the 0.3 mm line-type calibration slide measured from the bottom-view and side-view images captured with the distance between the mirror and the sample at 2 mm, 3 mm, and 4 mm. M) The optical path in the cell culture medium and the chromatic aberration generated. N) The fitting curve between *NA* and relative offset distance between 486.1 nm to 656.3 nm (Δ*l*) when *d* is 1 mm, 2 mm, 3 mm, and 4 mm. O) The bottom-view and side-view images of 0.03 mm microscope calibration slide captured with 4×, 10×, 20×, and 40× objective lenses in ddH_2_O and DMEM medium. P) The length of the 0.3 mm line-type calibration slide was measured in ddH_2_O and DMEM mediums through the bottom view and side view. (error bars = SD, n = 5)

Bottom-view images were captured using an inverted microscope, providing measurements like diameter and width of spheroids (*x*-*y* plane, **Fig. 1C**). Side-view images, facilitated by the mirror, offered additional measurements such as thickness, height, and contact angle (*x-z* plane, **Fig. 1C**). The principle of non-destructive *in situ* observing the samples from both bottom-view and side-view was illustrated in **Fig. 1D**. For bottom-view, the standard microscope setup sufficed. For side-view, light from the source, either reflected by the sample or transmitted through it, was redirected into the microscope’s objective by the first surface mirror. This placement enabled capturing side profiles (**supplementary information sVideo1**). **Fig. 1E** shows the experiment setting of observing the bottom-view and side-view of the samples. No modification to the microscope was done to achieve non-destructive *in situ* observation of the bottom-view and side-view of cell spheroids at a low cost.

Maintaining image authenticity is a prerequisite for using a first surface mirror to achieve side-view observation on an inverted microscope. Firstly, we tested image quality using a 0.3 mm round-type microscope calibration slide (ChenZheng Precision Tools, Suzhou, China). For bottom-view imaging, the sample was placed directly on the stage. For side-view imaging, it was cut and attached vertically to polyethylene foam double-sided tape (**Fig. 1F**). Images captured with 4×, 10×, and 20× objective lens showed no significant quality difference between bottom-view and side-view observation. However, it was noted that the 40× objective lens’ field has limitations in imaging (**Fig. 1G, 1H**).

To study the impact of the distance between the sample and the mirror on image acquisition, a 0.03 mm line-type microscope calibration slide was fixed vertically to a base at a 90° angle to the horizontal plane (**Fig. 1I**), and another microscope calibration slide to the bottom surface (**Fig. 1J**). **Fig. 1K** shows images captured between the mirror and the sample at 2 mm, 3 mm, and 4 mm distances. Side-view images remained similar across these distances. However, at 4 mm, the 40× objective couldn’t focus due to exceeding its working distance (3.6-2.8 mm). The results demonstrated that as the sample was within the working distance of the objective lens, there was no significant change in the apparent size of the sample observing with 4×, 10×, 20×, or 40× objective lens (**Fig. 1L**).

For capturing spheroid images, the first surface mirror was placed in a cell culture medium to maintain growth conditions. Typically, introducing different media into the light path causes refraction and potential chromatic aberration. For instance, light passing through the DMEM medium undergoes such refraction. To evaluate the impact of chromatic aberration on imaging resolution, we analyzed the offset values of the C-line (656.3 nm) and F-line (486.1 nm) using chromatic aberration analysis (**Fig. 1M**). The analysis follows:

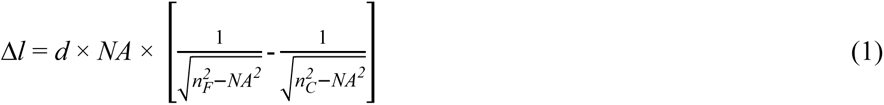

where *NA* is the numerical aperture, Δ*l* is the relative distance offset, *d* is the total length of the folded light path from the sample to the bottom of the petri dish, *n_F_* is the refractive index at 486.1 nm, and *n_C_* is the refractive index at 656.3 nm. Higher *NA* and longer *d* increase chromatic aberration, affecting observations (**Fig. 1N**). But, as observed in **Fig. 1O**, enabling the automatic white balance function during the imaging process can effectively minimize the chromatic aberration. **Fig. 1P** shows that images at 4× and 10× in ddH_2_O or DMEM had no significant differences compared to bottom-view images in air. Considering the distance between the sample and the mirror, and the culture medium-induced refraction, using the first-surface mirror for high magnification (40×) side-view imaging is not recommended.

### The side-view observation petri dish/device allows non-destructive, real-time observation of the bottom- and side-view of the 3D spheroid

Growing cells at agarose micro-wells or agarose-filling microplates is one of the effective methods to generate 3D spheroids (Caprio and Burdick, 2023; Froehlich et al., 2016). For imaging, our focus was on light emerging from the spheroid surface, not internal refraction or scattering. **Fig. 2A** illustrates that during bottom-view imaging, the agarose micro-well directly contacts the bottom surface, requiring the image to pass only through the agarose layer to reach the microscope objective. For side-view imaging (**Fig. 2B**), light passes sequentially through the agarose micro-well, culture medium (DMEM), reflects off a first-surface mirror, and finally enters the microscope objective. Therefore, the optical path through the agarose was the same for both bottom-view and side-view imaging (with both bottom and wall thicknesses being 1.5 mm), resulting in identical refraction introduced by the agarose micro-well in both viewing modes. Next, we studied the impact of agarose and culture medium on imaging quality. To support cell spheroid growth, we used micro-wells formed from 2 wt% agarose (Pan et al., 2023a). Absorption spectra of 1 wt% agarose, 2 wt% agarose, and DMEM culture medium (each 1 cm thick), were measured using a fiber optic spectrometer (PG2000-Pro-EX, Fuxiang Optics Co., Ltd., Shanghai, China), with ddH_2_O as the control (**Fig. 2C**). It was observed that both 1 wt% and 2 wt% agaroses partially absorb light in the 400-600 nm range, while DMEM medium, containing phenol red as an indicator, had absorption peak at 420 nm and 550 nm (Zhikhoreva et al., 2018). The spectra of the aforementioned samples under the CEL-TCX250 (Xenon lamp light source) are shown in **Fig. 2D**. The DMEM medium and agarose primarily absorbed light in the 400 nm to 600 nm range, with less absorption from 600 nm to 800 nm, causing the spheroids’ images to appear magenta. During imaging, we used automatic white balance correction to minimize the impact of color on the cell images. Since this study focused mainly on the morphology of the spheroids rather than their color information, the introduction of the culture medium and agarose did not have a significant negative impact on the experimental results.

**Fig. 2.**
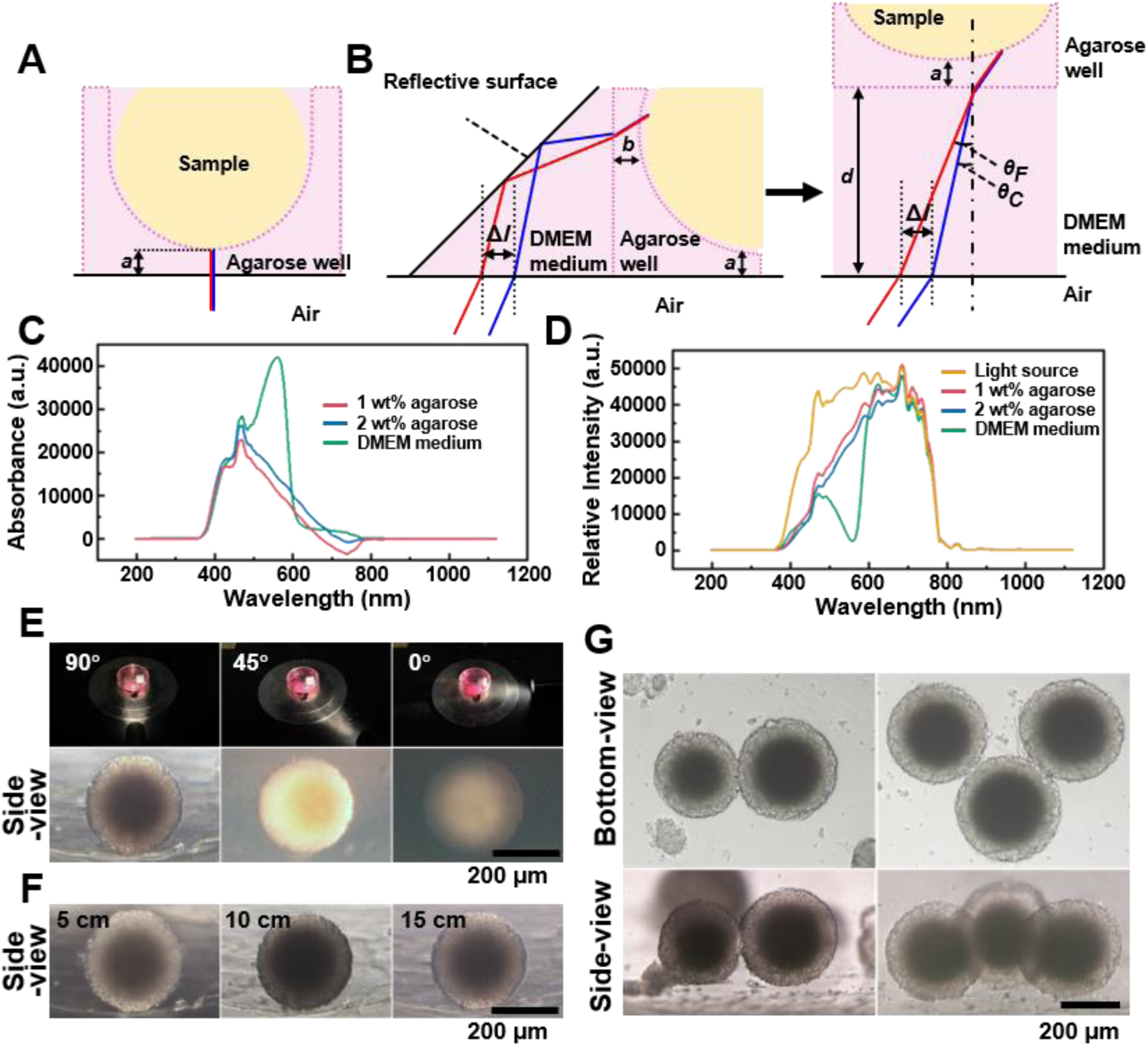
A) Light path of the bottom-view observing the spheroid in the agarose micro-well. B) Light path of side-view observing of the spheroid in the agarose micro-well. C) The absorption spectrum of the 1 wt% agarose, 2 wt% agarose, and DMEM culture medium. D) The spectra of the 1 wt% agarose, 2 wt% agarose, and DMEM culture medium under the CEL-TCX250 (Xenon lamp light source). E) Images quality display of spheroid under the 90°, 45° and 0° lighting. F) Images display of the spheroids under 5 cm, 10 cm, and 15 cm lighting. G) Using the side-view observation petri dish/device can be used to photograph multi-spheroids samples.

To optimize conditions for side-view imaging, we chose a xenon lamp for its broad wavelength coverage and adjustable angle. **Fig. 2E** shows that a 90° angle between the lamp and first-surface mirror produced the best details of the side-view images, while angles of 45° and 0° resulted in uneven and poorly lit images. Maintaining the 90° angle, we examined the effect of varying distances between the light source and sample (**Fig. 2F**). At 5 cm, internal structures (dark core) were clear, but uneven background illumination complicated edge distinction. At 10 cm, background uniformity improved, though overall illumination remained irregular, and the dark core was not resolved. At 15 cm, the sample background contrast was optimal, with clearly defined edges and discernible internal details, including the dark core. By adjusting the position of the sample stage, it became straightforward to locate and observe the side-view of spheroids within agarose micro-wells (**supplementary information sVideo2 and 3**). And the side-view observation petri dish/device can be used to photograph not only a single spheroid but also multi-spheroids samples (**Fig. 2G**). Collectively, for observing and imaging the side morphology of samples using the side-view observation petri dish/device, particularly for samples in culture medium, we recommended using a non-divergent light source with broad-spectrum coverage. This light source should be positioned approximately 15 cm directly behind the sample, forming a 90° angle with the mirror. Samples should kept as close to the mirror as possible, with the distance between the sample and the mirror kept within the working distance of the objective lens. Considering the diameter of 3D spheroids ranges from 200-500 μm, side-view images were captured using 4× or 10× objective lenses in subsequent studies.

### Tracking the growth dynamics of spheroids from multi-angle observation

Tracking the growth images of cell spheroids enables further determination of their volume or size, which is significant for sample screening. However, conventional methods rely heavily on bottom-view images because it is challenging to monitor the side-view changes *in situ*. This study tracked the spheroid formation process in the agarose micro-well from a seeding density of 1×10^4^ cells-per-well using the inverted microscope. Time-lapse images of the bottom-view and side-view of the cells within the agarose micro-well, as shown in **Fig. 3A**, record the morphological changes of the cells during spheroid formation. Initially, the newly added human prostate cancer cells DU 145 scattered in the micro-well. After 3 h of incubation, the cells clustered to form a ring structure. From the side-view, the thickness changes of the cell cluster were negligible, suggesting that the cells tend to aggregate at a similar planar. As the cells continued to aggregate, the cluster gradually became more compact and formed a round shape after 60 h of incubation. However, the side profile of the same spheroid shows that the thin layer of the cell disk gradually packed, and as the *x-y* dimension decreased, the height/thickness of the cell aggregate increased (**Fig. 3B**). We also placed the side-view observation petri dish/device on a live cell monitoring system (MoniCyteTM B-100, Jiangsu Rayme Biotechnology, China) to capture a time-lapse video of the side-profile changes spheroid formation process of 1×10^4^ DU 145 cells over 60 h (**supplementary information sVideo 4**). Through those time-lapse images/video, the width and height/thickness of the spheroid can be measured. As suggested in previous studies, when the height/thickness and width ratio of the cell aggregate approaches 1, it indicates that the spheroid structures have formed. In previous studies, spheroids had to be sealed or fixed in an agar block before the side-view morphology could be observed(Pan et al., 2023a). The side-view observation device allows for non-destructive tracking of the growth dynamics of the same spheroid in both bottom-view and side-view, which enables the depict the 3D structural characters of cell aggregates.

**Fig. 3.**
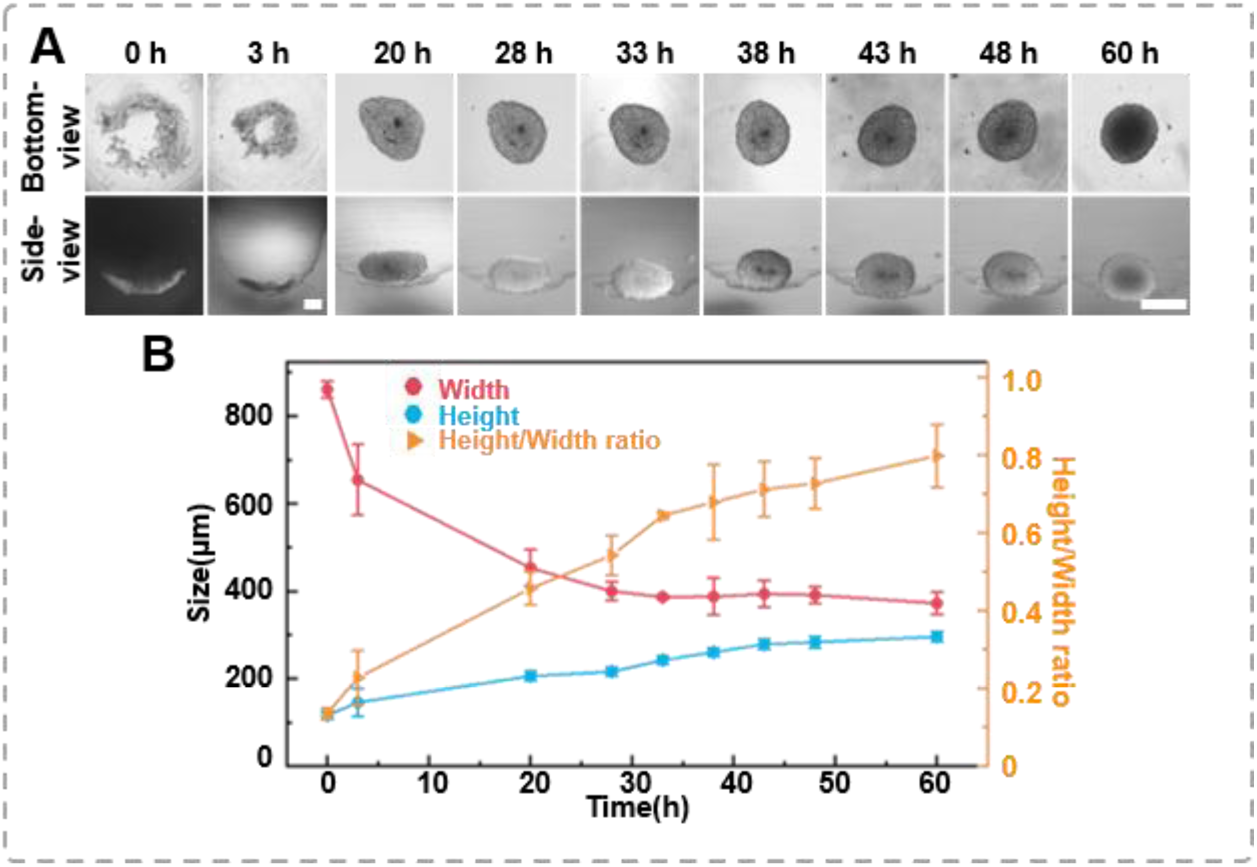
The formation dynamics of the spheroids. A) Time-lapse images of DU 145 spheroid formation in an agarose well (1×10^4^/well, scale bar = 100 μm); B) Changes in height and width during the growth process of the spheroids, as well as variations in height-to-width ratio.

### Characterization spheroids 3D structure changes during spheroid migration

Cell migration from or leaving tumor spheroids is crucial for the dissemination of solid tumors and the occurrence of secondary metastases, both of which are life-threatening. Previously, spheroids were placed on a substrate, and their migration capability was characterized by cell migration distance (Carvalho et al., 2022; Kim et al., 2022). This study placed tumor spheroids on a petri dish for a cell migration assay. As the bottom-view morphology shown in **Fig. 4A**, cells extruded from the spheroid, form ing sprouts. With prolonged culture time, cells from the spheroid periphery migrated outwards, forming a single-layer cell region surrounding the spheroid. The migration distance is the difference between the total width (spheroid + migrating length) and the inner width (spheroid only). As shown in **Fig. 4B**, the distance occupied by the migrating cells expanded from 0 μm to 2894.59±127.08 μm after 100 h of migration. It was important to note that the average inner width only decreased by 23.87 μm (a change rate of 6.8%).

**Fig. 4.**
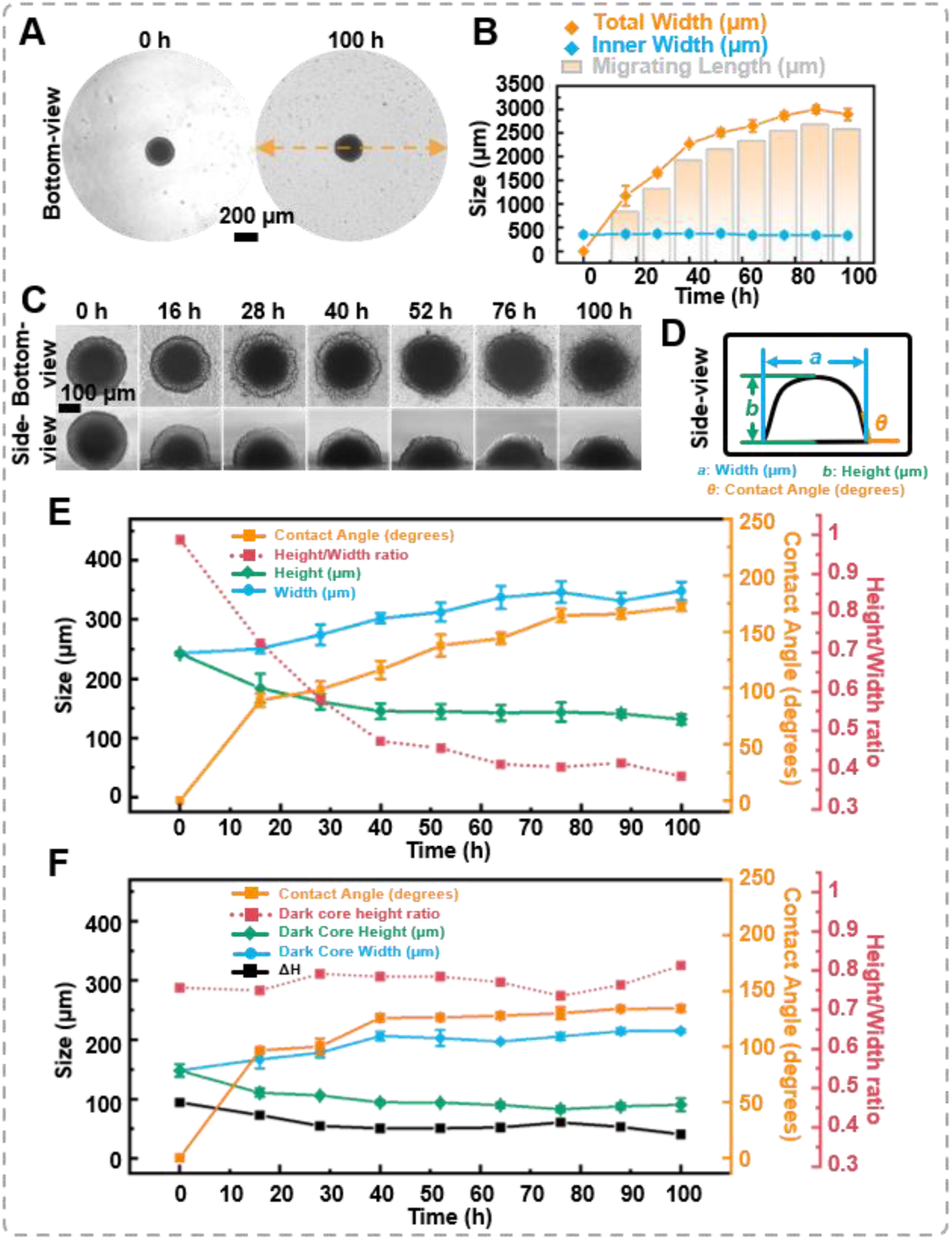
The migration of the spheroids. A) The bottom-view images of the spheroid in the petri dish, cells extrude from the spheroid, forming sprouts; B) Statistics on the changes in total width, inner width, and migration distance of spheroids during migration process; C) Time-lapse bottom-view and side-view images of DU 145 spheroids during migration process; D) Definition of the measured data points based on the side-view images of spheroids; E) Trend of morphological changes in the outer layer of spheroids during the migration process; F) Trend of morphological changes in the dark core of spheroids during the migration process. (error bars = SD, n = 3)

Simultaneously, with the assistance of the multi-angle observation device, variations in the spheroid’s height and contact angle during the migration process were recorded (**Fig. 4C and supplementary information sVideo 5**). **Fig. 4D** elucidates the measurement benchmarks for the side-view images. The contact angle between the petri dish surface and the spheroid was measured using the angle function of ImageJ. As shown in **Fig. 4E**, when the spheroid was placed on the petri dish, the thickness of the spheroid was 242.34±2.85 μm, and the contact angle between the surface and the spheroid was 0°. After 28 h of migration, the thickness of the spheroid gradually decreased to 161.11±13.08 μm, and the contact angle changed to 98.52±7.57°. Throughout 100 hours of migration, the thickness and the contact angle of the spheroids’ changed to 131.33±8.34 μm and 172.08±3.75°, respectively. Furthermore, from the side-view images, the ImageJ’s measurement function revealed that the dark core area was 46,125.40 μm² at 0 h and only decreased to 43,302.44 μm² at 100 h. Based on the side-view images, we calculated the difference between the height of the outer spheroid and the height of the internal dark core, denoted as ΔH. It was found that the height of the dark core decreased by 57.63 μm, while the total spheroid height decreased by 111.01 μm after 100 h of migration. The dark core height ratio increased from 0.61 to 0.69, suggesting that the proliferating cells at the outer region of the spheroid were the prominent participants in the migration process (**Fig. 4F**). Combining the bottom-view and side-view images, it was found that as the average width increases, the *z*-direction height of the spheroids decreases, leading to a gradual decrease in the height-to-width ratio of the cell spheroids and an increase in the contact angle of the cell spheroids with the petri dish.

### Association between the thickness of the spheroid and the appearance of the dark core of the 3D spheroid

A dark core within 3D cell spheroids is a phenomenon commonly observed. Biologically, the dark core is believed to represent necrotic or quiescent zones arising from limited diffusion of oxygen and nutrients within the spheroid. In addition, it is important to notice that, optically, the depth of light penetration in microscopy can be hindered by the spheroid’s thickness, leading to decreased signal intensity and diminished image quality from deeper regions. This effect is further amplified in larger spheroids by the increased number of cellular layers and a dense extracellular matrix, which scatter light and obscure core details. The side-view function of the proposed device allows tracing the formation of the visually distinguishable dark core to investigate the relationship between the dark core, spheroid size, and the optical limitations of light microscopy using DU 145 cell spheroids with different initial seeding densities (2.5×10^3^, 5×10^3^, 1 ×10^4^ and 2×10^4^ cells/well). The appearance of the dark core and its correlation with spheroid thickness were analyzed using the bottom-view and side-view of the spheroids.

As **Fig. 5A** shows, it was found that the time to dark core formation was inversely related to the initial seeding density. For example, spheroids initiated with 2.5×10^3^ cells exhibited a dark core between 48-60 h, achieving an average *z*-direction growth rate of 4.18 μm/h. These spheroids reached a nearly spherical shape with an average height of 250.67 μm and a maximal height-to-width ratio of 0.99 by 60 h. With an increase in seeding density to 5.0×10^3^ cells, the dark core appeared earlier (38-43 h), and the spheroids achieved a height-to-width ratio of 0.82 by 43 h and approached a spherical shape by 60 h, with a width between 270-290 μm. At a seeding density of 1.0×10^4^ cells, the dark core developed between 28-33 h, with a *z*-direction growth rate of 7.35 μm/h. The spheroids presented a height-to-width ratio of 0.54 at 33 h, evolving into a flattened spherical shape by 60 h because the side-view images indicate an elongated, bean-shaped profile. The highest seeding density of 2.0×10^4^ cells resulted in the earliest dark core formation (20-28 h), with a z-direction growth rate of 9.43 μm/h. These spheroids displayed the most pronounced elongation, with a height-to-width ratio of 0.49 at 28 h, further flattening by 60 h. We analyzed the relationship between dark cores’ appearance and width using conventional bottom-view images of spheroids (**Fig. 5B**). It is evident that with fewer seeding densities, the width of the spheroids gradually decreases over time, while the appearance of the dark core occurs later. The purple square highlight indicates the time points when the dark core appeared. Based on the trend observed in the images, we can predict that at lower seeding densities, the time for the appearance of dark cores in the spheroids will be longer, and vice versa. However, irrespective of the seeding densities, the *z*-direction height of the spheroids at the onset of dark core formation remained relatively consistent, approximately 250±15 μm (the yellow-colored zone in **Fig. 5C**), suggesting that there is a correlation between the formation of dark cores in spheroids and the *z*-direction height of the spheroids.

**Fig. 5.**
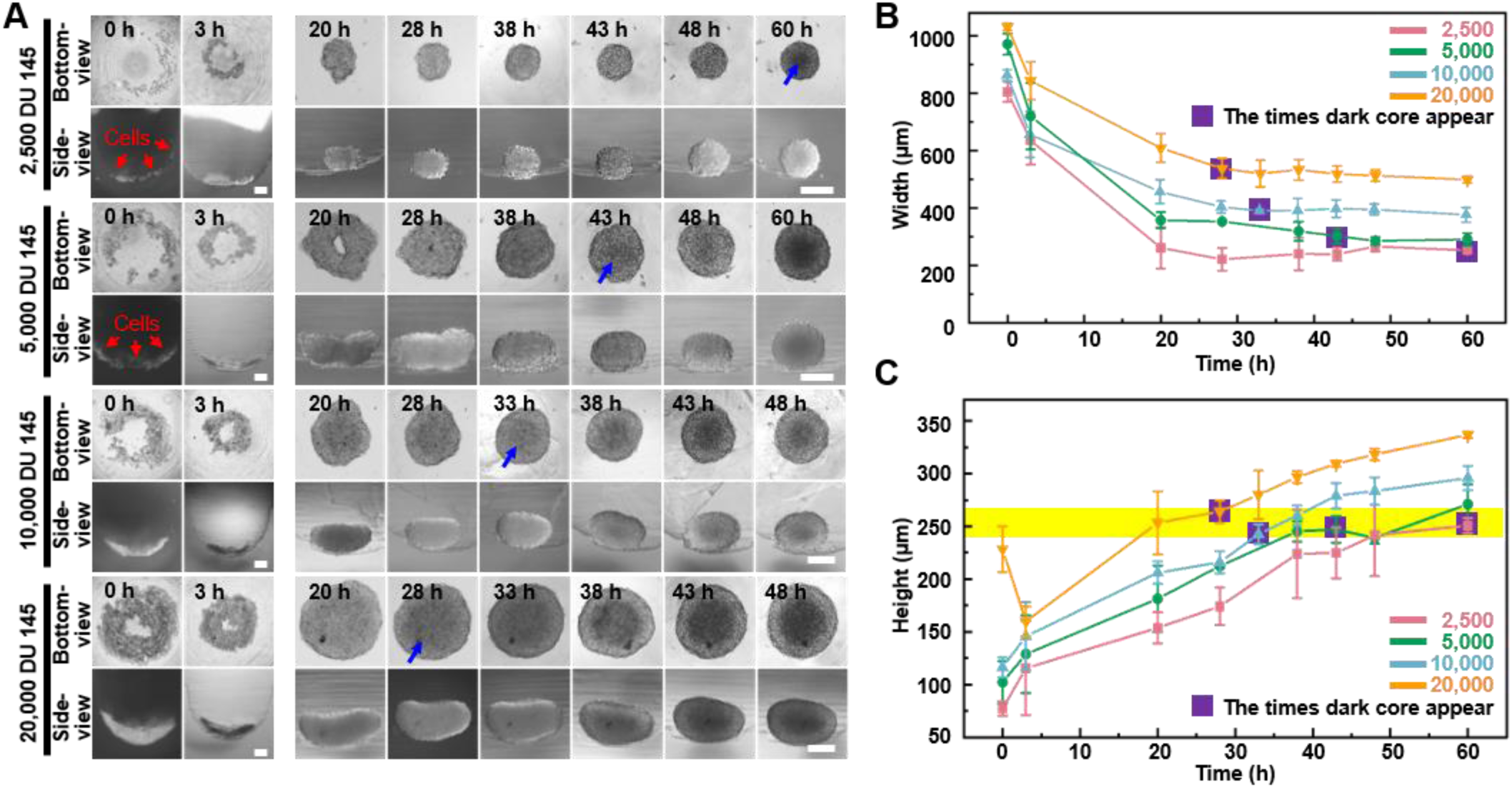
The dynamics of dark zone formation within the 3D spheroid. A) Time-lapse bottom-view and side-view images of DU 145 spheroids formed from different initial cell concentrations (2.5×10^3^, 5×10^3^, 1×10^4^ and 2×10^4^ cells/well). Blue arrows indicate the appearance of observable dark cores; B) Relationship between spheroid width changes and dark core appears at different initial cell concentrations analyzed through bottom-view images; C) Relationship between spheroid height changes and dark core appears at different initial cell concentrations analyzed through side-view images (the yellow stripe indicates the height of 250±15 μm). (error bars = SD, n = 3)

### Characterization spheroids 3D structure changes during spheroid fusion

Based on observing the spheroids’ formation and migration, we hypothesize whether the multi-angle observation device facilitates the observation of more intricate cellular spheroids’ structural changes through side profile. Fusion is a crucial step for tissue development. In tissue engineering, 3D spheroids fusion is vital in fabricating micro-tissues (Gcj et al., 2021; Laschke and Menger, 2017). However, the fusion process of spheroids exhibits a highly random situation, making it challenging to maintain the fusion state while observing it *in situ* to reconstruct the structural changes during spheroid fusion. Herein, the side-view device was applied to conduct *in situ* observations of the fusion process of cell spheroids. Spheroids of 1×10^4^ DU 145 cells per well were cultured for 7 days in agarose micro-wells for fusion assay. As the bottom-view images shown in **Fig. 6A-i**, the two spheroids contact each other and gradually merge at the contact region. The contact angle, contact length, and doublet length value can all be retrieved from time-lapse images (**Fig. 6B**). First, the contact angle between spheroids increased from 30° to 45°. Next, the two spheroids’ contact length (neck) increased with co-culture time, with the most significant change occurring within the first 3 h. The doublet length of the fusion formed by two spheroid decreased by 22.5% from 453.87 μm to 367.19 μm after 48 h of fusion. Though the bottom-view depicts the *x*-*y* dimension changes of the spheroids during the fusion process, predicting the phenome occurring on the side profile of the spheroid is challenging, especially for those cell aggregates without ideal spherical structures (the red arrow denoted in **Fig. 6A-ii**). With the assistance of the side-view device, it was found that the two spheroids contacted and formed symmetric necks that gradually fused (**Fig. 6A-i**). As shown in **Fig. 6B**, the contact length between two spheroids progressively approaches the *z*-direction thickness, and the contact angle between the two spheroids reached 171.10±13.94° after 100 h of fusion. The green box in **Fig. 6B** highlights that the fusion process of two spheroids can be transformed into a process where the contact length approaches the height. Thus, supplemented by the side-view perspective, modeling, and reconstruction of the cell spheroid fusion process become feasible (**Fig. 6C**). The vertical lines in the figure indicate the position where the arc representing the contact angle at the fusion site of the cell spheroids tangentially intersects both spheroids. A wider width between the two vertical lines indicates a higher fusion process between the two spheroids. Additionally, we observed an intriguing phenomenon: if the initial cell spheroid morphology is not entirely regular or spherical (the red arrow denoted in **Fig. 6A-ii**), the side-view profiles enable to track the changes in the particular region except the fusion region. Therefore, the reconstruction from the side-view perspective can be more comprehensive.

**Fig. 6.**
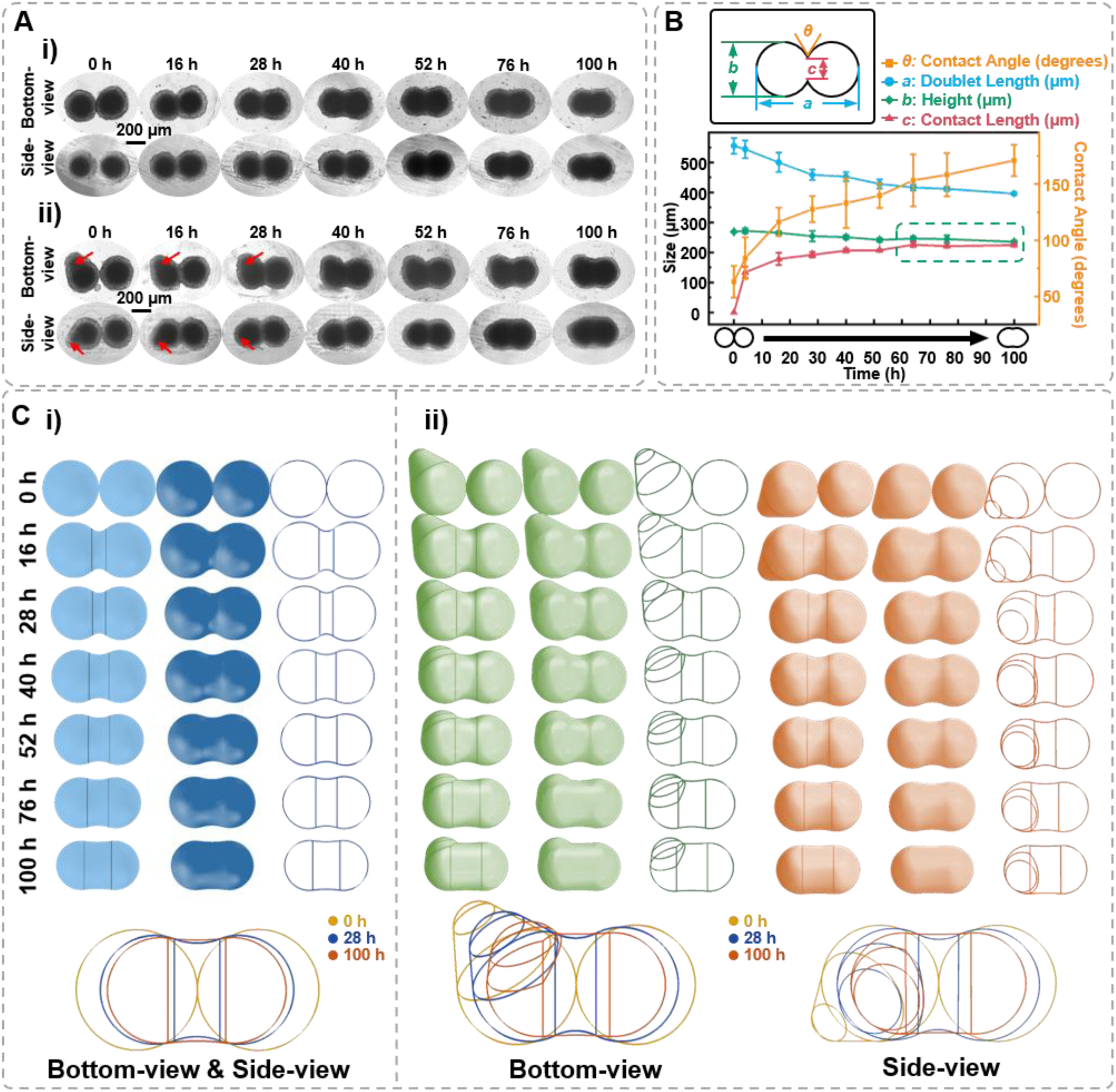
Characterization of the spheroids fusion process. A) Time-lapse bottom-view and side-view images of fusion body formed by 7-day-age DU 145 cell spheroids with ideal ball-like structure (i) and irregular cell spheroid (ii). Red arrows point to the protruding part of the cell spheroid. B) Definition of the measured data points based on the side-view images of fusion spheroids and the changes in doublet length, height, contact angle, and contact length during the fusion process. C) Three-dimensional spheroid fusion process modeling using bottom-view and side-view images of ideal ball-like structure (i) and irregular cell spheroid (ii). The vertical lines on the model surface illustrate the position where the arc tangentially intersects both spheroids. (error bars = SD, n = 3)

### Evaluate the impact of anti-tumor therapeutic reagents on the spheroid integrity

Spheroids are important in vitro cell models for drug testing, and the impact of drug effects is typically evaluated by the morphology and size of the spheroids (Lu et al., 2015). In this study, we investigated the effects of doxorubicin (DOX), one of the major chemotherapy reagents currently clinically used, and natural killer (NK) cells, a type of cytotoxic lymphocyte, on the 3D structure of DU 145 prostate cancer spheroids.

First, DU 145 spheroid (1×10^4^/spheroid) was cultured for 3 and 7 days and treated with 50 μg/mL DOX for 96 h. The spheroids aged for 3 days exhibited a relatively smaller dark core, with a diameter of 100.67±24.65 μm, and the dark core of 7 days spheroids is 168.32±5.98 μm. Time-lapse imaging (**Fig. 7A**) revealed that with extended DOX treatment, the outer region of the 3-day-old spheroids gradually lost its smoothness, as observed in both bottom-view and side-view. Furthermore, the visually dark core area was significantly enlarged. Similar morphological changes were noted in the 7-day-old spheroids. Meantime, it was found that DOX affects the height and width of the spheroids, causing them to expand within a certain period and then decrease in size as time progresses (**Fig. 7B**). It is worth noting that despite the significantly changed surface smoothness, the ratio of overall height-to-width remained largely unchanged, suggesting the spheroid’s overall spherical structure was preserved.

**Fig. 7.**
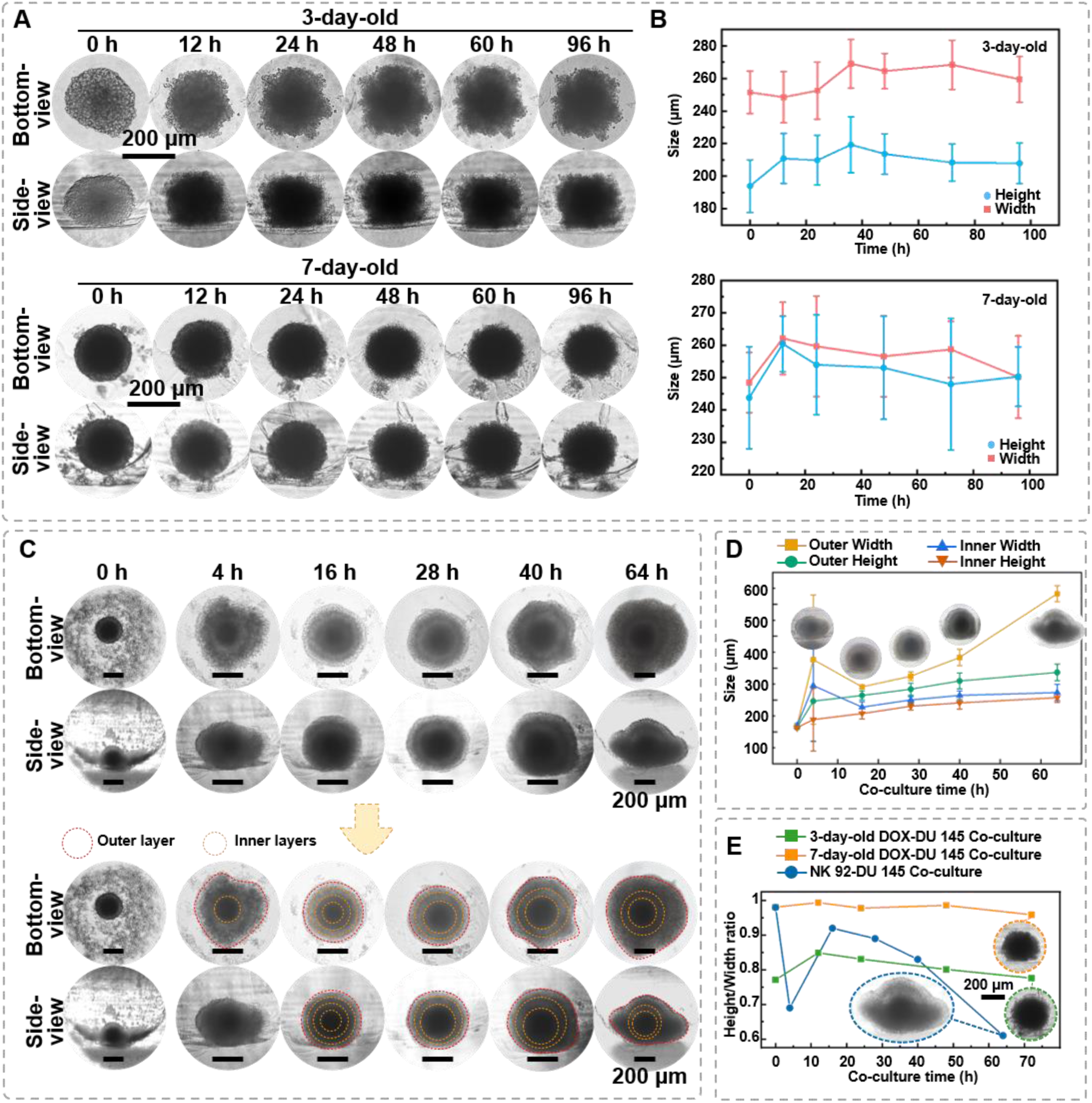
The *in situ* observation of DU 145 spheroids under the killing effect of DOX and NK cells. A) Time-lapse bottom-view and side-view images of 3-day-age and 7-day-age DU145 cell spheroids (1×10^4^/spheroid) under DOX (50 μg/mL) treatment. B) The changes of DU 145 spheroids’ width and height upon DOX treatment; C) Time-lapse bottom-view and side-view images of 7-day-old DU145 cell spheroids (1×10^4^/spheroid) co-cultured with 1×10^4^ NK-92 cells; D) Changes in internal and external width and height of 7-day-old DU145 cell spheroids (1×10^4^/spheroid) co-cultured with NK-92 cells; E) The different performances of DOX and NK cells killing on the overall height-to-width ratio of the spheroids. (n = 3, error bars = SD)

Next, we also explored the role of NK cells, a type of cytotoxic lymphocyte critical in the immune system’s defense against tumors, as model immune cells. As depicted in **Fig. 7C**, the 1×10^4^ NK cells spread in the micro-well at 0 h of the co-culture. After 4 h of co-culture, NK cells gathered around and enveloped the DU 145 spheroid as increased diamension in both bottom and side profiles. After 16 h of co-culture, unlike the distinct boundary between dark core and translucent zone, referring to necrotic and proliferating regions of the spheroid, observed before co-culture, four optically different regions could be distinguished from both bottom-view and side-view, suggesting that NK cells infiltrated the DU 145 spheroids from all directions. At this time point, the overall height and width of the spheroid were 314.47±13.85 μm and 341.20±5.81 μm, respectively. Thus, the height-to-width ratio was close to 1, suggesting a spherical structure. After 28 h of co-culture, only three optically different regions could be distinguished, and the distinction between the inner two regions became less obvious than at 16 h. Moreover, the side-view images found that the spheroid appeared to collapse as the height-to-width ratio decreased to 0.89. However, measurements indicated expansion in the *x-y* and *x-z* directions, with overall growth rates of 9.78% and 6.22%, respectively. After 40 h of co-culture, the collapse of the outer layer was apparent, and the height-to-width ratio was 0.83. At the same time, the spheroid’s overall size continued to increase, with growth rates of 15.74% and 7.73% in the *x-y* and *x-z* directions, respectively. After 64 h, the outer cell layer had completely collapsed. Cells had detached onto the bottom of the agarose well, losing the spherical shape (**Fig. 7D**). By correlating the size growth rates observed in side-views with corresponding time points, the significant increase in the width at 4 h was attributed to the external aggregation of NK cells on the spheroid. Over time, NK cells gradually envelop the spheroid without a noticeable change in the *z*-direction. From 20 to 60 h, both bottom- and side-directions exhibited growth until the upper region collapsed downwards at 64 h.

Analyzing the 3D structural changes of spheroids under DOX and NK cell treatment indicates that through different mechanisms, antitumor agents result in distinct morphological features of the spheroid. **Fig. 7E** compares the height-to-width ratio of the spheroids treated with DOX and NK cells for 64 h. The cytotoxic effect of NK cells on spheroids resulted in a 37% decrease in the height-to-width ratio, while there was only a 0.64% change for 3-day-old spheroids treated with DOX and a 2.24% change for 7-day-old spheroids treated with DOX. Therefore, viewing the spheroids from different angles, such as bottom- and side-view, would provide precise structural changes in the drug-induced killing effects.

## Discussion

We developed a multi-angle observation device using 3D printed components and a first surface mirror, which costs less than $1 to produce, to address the current limitations in observing 3D cell spheroids. First, the proposed side-view observation device was validated for observing the spheroid formation and migration process from both bottom- and side-view. As shown in **Fig. 3**, side-view observation provides detailed insights into the spheroid formation, including the 3D structure of the spheroid and its height at a given time point. Past studies have often assumed spheroids to be centrally symmetrical (Filippo Piccinini et al., 2015), elliptical(James D Murphy et al., 2011), or spherical(Senavirathna et al., 2013). It is evident that as the spheroid grows, its morphology does not directly conform to a regular shape (**Fig. 3A and B**). *In situ* tracking and imaging from a side-view perspective can facilitate modeling to reconstruct the real shape of the spheroid accurately and enable researchers to gain a more comprehensive understanding of the 3D structure and growth kinetics of spheroids without disrupting their natural development.

This device offers a unique opportunity to study the 3D aspects of cell migration from tumor spheroids (**Fig. 4C**). The side-view images allow us to analyze the interaction between the spheroids and substrate by measuring the contact angle. Moreover, the changes in the dimensions and area of the dark core were less intense than the overall changes of the spheroid, suggesting that the proliferating cells at the outer region of the spheroid were the main participants in the migration process. To the best of our knowledge, this is the first time the side profile changes of the spheroid during a migration assay have been depicted. By monitoring both bottom and side changes, researchers can gain valuable insights into the spatial distribution of migrating cells and the dynamics of the migration process to elucidate the mechanisms underlying tumor cell invasion and metastasis, potentially leading to the development of new therapeutic strategies targeting these processes.

The observation of the dark core in the spheroid highlights the presence of a necrotic or quiescent region, which is a common feature of larger tumor spheroids(Browning et al., 2021; Gcj et al., 2021; Laschke and Menger, 2017). As demonstrated in **Fig. 5**, we observed that spheroids formed at different cell seeding densities, regardless of whether they approached a height-to-width ratio close to 1, began to develop a dark core at an overall height of approximately 250±15 μm. This consistency suggests that the timeframe for dark core formation is primarily determined by the increasing *z*-direction height, with a critical threshold of approximately 250 μm may indicate the limit of effective oxygen diffusion, beyond which cellular functionality becomes compromised. The observation aligns with previous findings that suggest a maximum viable spheroid diameter of 500 μm, beyond which central necrosis becomes inevitable (Pan et al., 2023a). This clear observation was not achievable using only bottom-view images, emphasizing the importance of real-time monitoring and 3D structural analysis of spheroids through side-view observation. For the spheroids’ fusion process, conventional observation using only bottom-view imaging might miss the critical structural changes on the other side of the fusion process (**Fig. 6A**). However, with the multi-angle observation device, we could assist in the observation of three-dimensional structural changes from the side, thus *in situ* monitoring of irregularly shaped spheroids, tissues, and organoids from multiple angles become possible.

Finally, we compared the cytotoxic effects of two different approaches, DOX and NK cells, on spheroids using the multi-angle observation device. **Fig. 7** shows the distinct outcomes of DOX treatment versus NK cell-mediated cytotoxicity: DOX treatment does not significantly alter the three-dimensional morphology of the spheroids, whereas NK cell-mediated cytotoxicity notably affects the overall height-to-width ratio of the spheroids. DOX exerts its effects by disrupting DNA and RNA synthesis, inducing ROS-related apoptosis, and triggering cellular senescence, among other mechanisms (D.Agudelo et al., 2016; Kong et al., 2022). These actions lead to morphological changes such as nuclear condensation, fragmentation, and cell shrinkage. The observed morphological features from both the bottom-view and side-view suggest that DOX may induce a form of cell death in tumor spheroids without disrupting their overall 3D structure (**Fig. 7A**). On the other hand, as shown in **Fig. 7C and D**, NK cells gathered around the tumor spheroids and gradually infiltrated into the spheroids, leading to the expansion of the overall size of spheroids. Moreover, the co-culture of NK cells and tumor spheroids resulted in significant structural changes, as the height-to-width ratio decreased from 0.98 to 0.61 after 100 h of co-culture. Considering that NK cells exert cytotoxicity by forming pores in the target cell membrane, leading to membrane disruption and cell lysis, the side-view profile indeed captured the gradually demolished spheroid’s 3D structure. Relying solely on bottom-view observations of co-cultured spheroids may not adequately predict side profile changes or the overall 3D structure, potentially resulting in volume prediction inaccuracies. Therefore, by accurately observing the responses of spheroids to different cytotoxic agents, researchers can design more precise drug induction protocols and develop more effective strategies for tumor eradication.

The proposed device is compatible with inverted microscopes, allowing for *in situ* bottom-view and side-view observation of 3D cell spheroids, a capability not previously reported. Compared to existing side-view microfluidic devices(Lee et al., 2020), our device offers lower production costs and is easier to manufacture. Additionally, the device features magnetic connections for easy positioning adjustment to observe multiple samples (Supplementary sVideo 4, and 5) and convenient disassembly for reuse and sterilization purposes. To further improve the capability of the side-view device to analyze the sophisticated detail of 3D spheroids, such as the visually optical distinct layers in NK-tumor spheroids co-culture (**Fig. 7D**), we are considering two avenues for optimization: improving the image capture process and enhancing image quality: one approach involves adding lenses in front of the first surface mirror. At the same time, the other entails designing complementary light sources to reduce light scattering within the spheroids during imaging. Once the image capture process is further enhanced, we can leverage machine learning to analyze the side-view morphology of the spheroids and organoids to predict their future growth patterns, potentially saving considerable screening time in preclinical drug testing.

## Materials and Methods

### Reagents

Agarose (for biochemistry, MW= 630.5, melting point 62-68 °C, PubChem CID 11966311) was purchased from Aladdin, China. Phosphate-buffered saline (PBS, pH 7.4) were purchased from Beijing DingGuo ChangSheng Biotechnology Co. LTD. Human prostate cancer cells DU 145 were obtained from Precella Inc. (Wuhan, China). The tumor cells were cultured in Dulbecco’s TM®Modified Eagle Medium (DMEM, Gibco, USA), containing 10% fetal bovine serum (Bio- channel, Nanjing, China), penicillin (100 U/mL), and streptomycin (100 μg/mL) at 37 ℃ in a 5% CO_2_ atmosphere. The NK-92 cells were cultured in NK-92 special medium Cat# CM-0530 (Precella, Wuhan, China).

All the chemicals were used as received and prepared in ultrapure water (PURELAB flex system, ELGA Corporation, IL, USA) without further purification. We employed a CEL-TCX250 xenon lamp light source from China Education Au Light Co., Ltd., Beijing, China, for side-view imaging. The microscope’s built-in light source was used for all other imaging.

### Design and fabrication of the multi-angle observation petri dish /device

The overall size of the device, as shown in **Fig. 8A**, is approximately 17 mm in height, with a petri dish diameter of 35 mm. Inside the 3D-printed handle and the frame, two cylindrical magnets with a diameter of 3 mm and height of 3 mm were embedded (**Fig. 8B**). To compare with the 35 mm petri dish, the height, width, and thickness of the first surface mirror are 20 mm, 4.5 mm, and 2 mm (**Fig. 8C**). A MoonRay UV DLP printer (SprintRay Inc., CA, USA) was used with photosensitive resin (CLEAR SP-RH1004) to fabricate all the 3D-printed parts with a speed of 50 mm/s, the height of each slice is 200 μm, and infill density is 50%. Before coming into contact with cell samples, the device is first cleaned overall using 75% alcohol and then sterilized by UV light irradiation for 30 minutes. The total cost of manufacturing this device is less than $1.

**Fig. 8.**
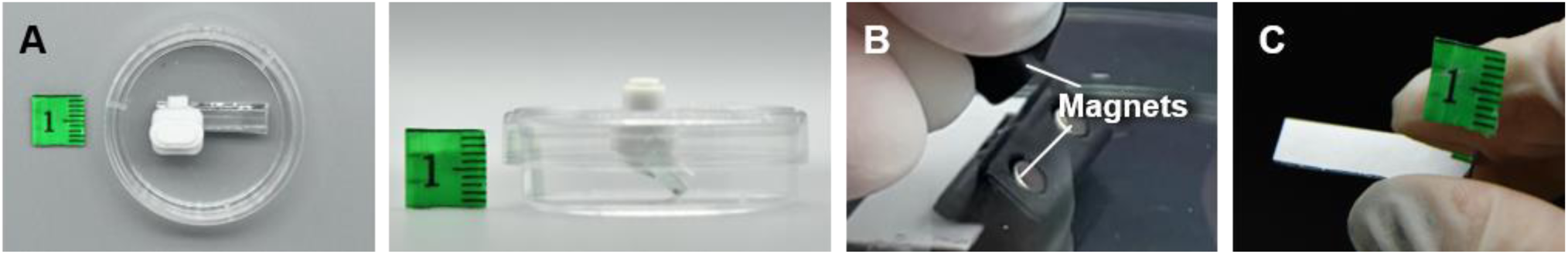
The display of the multi-angle observation petri dish/device. A) The top-view and side-view images of the multi-angle observation petri dish/device; B) The magnets inside the device.

### Measuring the spectra of the DMEM culture medium and agarose

The optical property of cell culture medium and agarose well for spheroids culture were characterized by fiber optic spectrometer. The opitc fiber was mounted on an optical platform, with one end connected to PG2000-Pro-EX fiber optic spectrometer (Fuxiang Optics Co., Ltd., Shanghai, China) and the other to a computer for spectral measurements. The light source (CEL-TCX250 Xenon lamp, light source) was placed at the same height as the fiber connection on the optical platform and preheated for 30 minutes. The 1 wt% and a 2 wt% agarose solution were prepared in advance, with 2 mL of each solution added to cuvettes and allowed to solidify for 15 minutes. During the absorption measurement, ddH_2_O was first used to calibrate the control sample under the light source. Subsequently, the 1 wt% and 2 wt% agarose block samples, as well as the DMEM medium, were measured sequentially. When directly measuring the spectra of the light source and the samples under the light source, the spectrum of the light source was measured first, followed by the 1 wt% and 2 wt% agarose gel samples, and finally the DMEM medium.

### Experimental setting of the multi-angel observation of the 3D spheroids

To capture the side-view images of samples under an inverted microscope, we used the Nikon ECLIPSE *Ti* microscope (TS100, Nikon, Japan). The objections we equipped are 4×, 10×, 20×, and 40× F objective lens (Nikon, Japan) with *NA* of 0.13, 0.30, 0.45, and 0.60, respectively. The corresponding working distance of 4×, 10×, 20×, and 40× F objective lens are 16.5, 15.2, 8.2-6.9, and 3.6-2.8 mm, respectively. The light sources we used to capture the bottom-view was D-LH/LC lamphouse (Nikon, Japan) and the light sources we used to capture the side-view was CEL-TCX250 (Xenon lamp light source, Beijing zhongjiao Jinyuan Technology Co., Ltd., China).

### Tracking the spheroids formation

The DU 145 cells were cultivated in agarose micro-wells using a previously reported procedure(Pan et al., 2023a). First, the agarose micro-well array was replicated from a 3D-printed mold. In brief, the replication process involved adding 700 μL of a 2% agarose solution to cover the 3D-printed mold, which was then allowed to solidify at room temperature (25°C). Once solidified, the agarose-based material was removed from the mold, resulting in an agarose micro-well array with a depth of 2 mm and a radius of 1 mm. The agarose micro-wells were then placed in the 35 cm petri dish. For tumor cell culture on the agarose micro-wells, the suspension with 2.5×10^3^, 5×10^3^, 1×10^4^ and 2×10^4^ DU 145-cells was added to each well. Subsequently, 600 µL of cell culture medium was added to cover the agarose base. The petri dish was placed in a cell incubator at 37 °C with 5% CO_2_. The bottom and side morphology of the spheroid formation was observed every 5 hours.

### Examining the spheroid migration/invasion from the sideview

1×10^4^ DU 145 cells/well in agarose micro-wells grown for 7 days were collected for migration assay. The spheroids were transferred into the petri dish with a multi-angle observation device installed on the lid. Then, the mirror was moved near the spheroid for observation using the handle on the lid. The petri dish was placed in a cell incubator at 37 ℃ with 5% CO_2_. The bottom and side morphology of the spheroid migration was observed every 12 hours.

### Observing the fusion of spheroids from different angles

1×10^4^ DU 145 cells/well in agarose micro-wells grown for 7 days were collected for migration assay. In brief, spheroids were pipetted to the agarose micro-well which was placed in the cell culture petri dish. The mirror was attached to the agarose micro-wells by adjusting the handle at the lid of the petri dish. The petri dish was placed in a cell incubator at 37 ℃ with 5% CO_2_. The bottom and side morphology of the spheroid fusion process was observed over 48 hours.

#### Evaluating the impact of the drug on the spheroid 3D structure

1×10^4^ DU 145 cells/well in agarose micro-wells grown for 3 and 7 days were collected for drug and NK-92 cells assay. In brief, spheroids were pipetted to the agarose micro-well which was placed in the cell culture petri dish.

***DOX***: DOX was diluted with cell culture medium to 50 μg/mL and added to the micro-wells seeded with DU 145 spheroids. Then the bottom and side morphology of the spheroid fusion process was observed over 48 hours at 37 °C with 5% CO_2_.

***NK-92 cells***: NK-92 cells were cultured in NK-92 special medium. For the co-culture assay, the NK-92 cells suspension was added to the micro-wells seed with DU 145 spheroids. The petri dish was placed in a cell incubator at 37 ℃ with 5% CO_2_. The bottom and side morphology of the spheroid fusion process was observed over 48 hours.

### Statistics analysis

Data are expressed as mean ± SD (standard deviation) of the number of biological replicates indicated in each figure legend. All the data analysis, bar graphs, and fitting curves were plotted by Origin (OriginLab Corporation, 2023, USA).

## Supplementary information

sVideo 1: Observation of the bottom-view and side-view of a tumor spheroid using the multi-angle observation device.

sVideo 2: Locating the side-view of the spheroid in the agarose micro-well through the bottom-view.

sVideo 3: Recording the side-view images of different spheroids in the agarose micro-well.

sVideo 4: Non-destructive *in situ* observation of the thickness changes during DU 145 cell spheroids formation process.

sVideo 5: Non-destructive *in situ* observation of the thickness changes during DU 145 cell spheroids migration process.

## Acknowledgments

This work was financially supported by the National Natural Science Foundation of China (No. 32171401), the Natural Science Foundation of Chongqing (CSTB2022NSCQ-MSX0808).

